# Discovery of tREP-18, a novel class of *t*RNA encoded peptide with potent leishmanicidal activity

**DOI:** 10.1101/2020.10.04.325217

**Authors:** Amrita Chakrabarti, Monika Kaushik, Juveria Khan, Deepanshu Soota, P Kalairasan, Sunil Saini, Siddharth Manvati, Jhalak Singhal, Anand Ranganathan, Soumya Pati, Pawan K. Dhar, Shailja Singh

**Affiliations:** Signaling Biology lab, Special Centre for Molecular Medicine, Jawaharlal Nehru University, New Delhi, India 110067; Synthetic Biology lab, School of Biotechnology, Jawaharlal Nehru University, New Delhi, India 110067; National Centre for Biological Sciences, Bangalore, India; Department of Life Sciences, Shiv Nadar University, Greater Noida UP, India 201 314

## Abstract

In the post genomic era, tRNA-derived fragments have emerged as a new class of non-coding gene regulators, those play crucial roles both at the transcriptional and translational levels, in different cellular biogenesis. However, none of the studies has ever asked whether tRNAs can also be translated into peptides with any biological significance. Thus, we present a novel hypothesis which suggested that; design and synthesis of tRNA-derived peptides from prokaryotic genome can be exploited for developing unique chemotherapeutics against neglected tropical diseases, like Visceral leishmaniasis (VL) and its aggressive form known as post kalazar dermal leishmaniasis (PKDL). To achieve this aim, we have used a novel system biology-based strategy, which involved; i) *mining of unique tRNAs from E. coli genome and their translation into peptide in silico*, ii) *designing of theoretical 3D models to evaluate their stability*, iii) *prediction of their biological activity by screening against anti-parasitic database to filter the lead peptide*. Based on this strategy, a unique tRNA-derived peptide (**tREP-18**) was selected, chemically synthesized, and used *in vitro* for elucidating its therapeutic significance against *L. donovani*, a causative agent of VL and PKDL. Our findings demonstrated that, **tREP-18** can impose high level toxicity to *L. donovani* promastigotes, by disrupting the ultrastructural cellular architect, destabilizing the mitochondrial membrane potential (ΔΨm), thus leading to drastic reduction in cell viability and proliferation. It also imparted high level of toxicity to BS12 a clinical isolate of PKDL. Conceivably, we for the first time reports a novel tRNA-derived peptide “**tREP18**” with excellent anti-leishmanial property, which can be further utilized for developing it as antileishmanial drug.

**One Sentence Summary:** Here we report a novel first-in-class tRNA encoded peptide and its super anti-leishmanial characteristics.

## Introduction

Transfer RNAs (tRNAs) are small non-coding RNAs (76-90 nucleotides in length) pivotal in bridging the DNA and the protein space^1^. By ferrying amino acids to the ribosomal interface, they bond amino acids in a specific sequence resulting in the formation of a polypeptide. Though well characterized and studied, the origin, evolution and translation of tRNA are still open questions which remain unanswered. Models have been proposed to imply direct duplication and evolution of RNA hairpin encoding gene^2^ and co-evolution of primordial tRNA with their association to translation machinery^3^. Furthermore, disrupted tRNA genes have been reported in the form of intron-containing tRNA^4^, split tRNA^5 6^, permuted tRNA in archae^7^. Evolutionary reasons behind these unexpected forms of tRNA gene sequences are still unclear^8^. Furthermore, asymmetric combinations of tRNA halves have been postulated to generate tRNA diversity^9^. Recent evidences showed that tRNA-derived small RNAs (tsRNAs) are generated following cleavage at specific sites by distinct nucleases. These tsRNAs have demonstrated multiple biological functions including stress-mediated signaling and as regulation of gene expression^10^. Additionally, they have been involved in RNA processing, cell proliferation, translation suppression, modulation of DNA damage response. These tsRNAs have also shown specific disease associations, including infectious diseases and metabolic abnormalities and neurodegeneration^10^.

However, no study has ever exploited the translation of tRNAs and/or the biological significance of tRNA-derived peptides. To address these unsolved puzzles, and exploit their therapeutic implication, we decided to design novel tRNA-based peptides and screen the same against visceral leishmaniasis (VL) the second largest neglected tropical diseases, endemic in several parts of the tropics, subtropics of Asia and Africa, and southern Europe, as reported by WHO (http://www.who.int/leishmaniasis/en/.2018.) This disease is caused by the species of protozoan obligate parasite, genus *Leishmania* (Kinetoplastida, Trypanosomatidae) which is usually anthroponotic in origin and transmitted by the bite of female phlebotomine sand flies. Leishmaniasis is exhibited in several forms among which the lethal ones are visceral and cutaneous one. Visceral Leishmaniasis (VL) is commonly known as Kala-azar, caused by the members of *Leishmania donovani*, and is prevalent in specific parts of India, Sudan and Nepal. About 3 million people are within the risk zone of infection and ∼200,000 to 400,000 new cases are reported each year. (http://www.who.int/leishmaniasis/en/) Among those patients who are apparently cured of Visceral Leishmaniasis (VL), can fall as prey to the hands of Post Kala-azar Dermal Leishmaniasis (PKDL). These patients are the strongest contender for being the disease reservoir. PKDL can manifest as life-threatening and/or disfiguring lesions ranging from innocuous self-healing cutaneous lesions to fatal visceralization of organs or dermal dissemination^11^. Thus, we have established a novel combinatorial approach involving *in-silico* analysis tools and synthetic biology application to characterize prokaryotic tRNA variants from mg1655 strain of *E*.*coli*. Based on *in silico* biology and structural analyses, we finally synthesized **tREP18**, as the lead peptide following screening against anti-parasitic database. Experimental evaluation revealed that, **tREP18** can cause sever morphological aberrations, destroy the membrane topology, distort ΔΨm, leading to complete abrogation cell viability and proliferation, and highly toxic to the BS12, an Indian clinical strain representing PKDL. This work reports a novel tRNA-derived peptide with enormous anti-leishmanial attribute.

## Results

### System biology-based analysis revealed tREP18 as a novel anti-parasitic peptide

We retrieved a total of 89 t-RNAs which were decoded from standard amino acids present in mg1655 strain of E. coli using Genomic t-RNA database. Out of this list, 29 peptides didn’t have any stop codons, thus considered for structure-based prediction. All the peptide sequences were successfully modelled, and total energy was calculated after energy minimization (Supp. Table 1). Out of 29, 9 peptides were filtered based on total energy with E-value, < 25, (selection criterion I, Fig 1) and considered for further analysis. The screening against the anti-parasite peptide databases exposed two peptides out of 9 with total energy E-value lower than 2 (selection criterion 2, Fig 1b Table 1). Among these two peptides, **tREP18** was selected as the lead compound due to its shorter length and E value 1.9 against antiparasitic database (Fig 1b Table 1, selection criterion 3). The scrambled peptides against the t-RNA-20 was modelled with two mutations in the sequence which leads to loss of secondary structure (Fig 1c).

**Figure 1:**
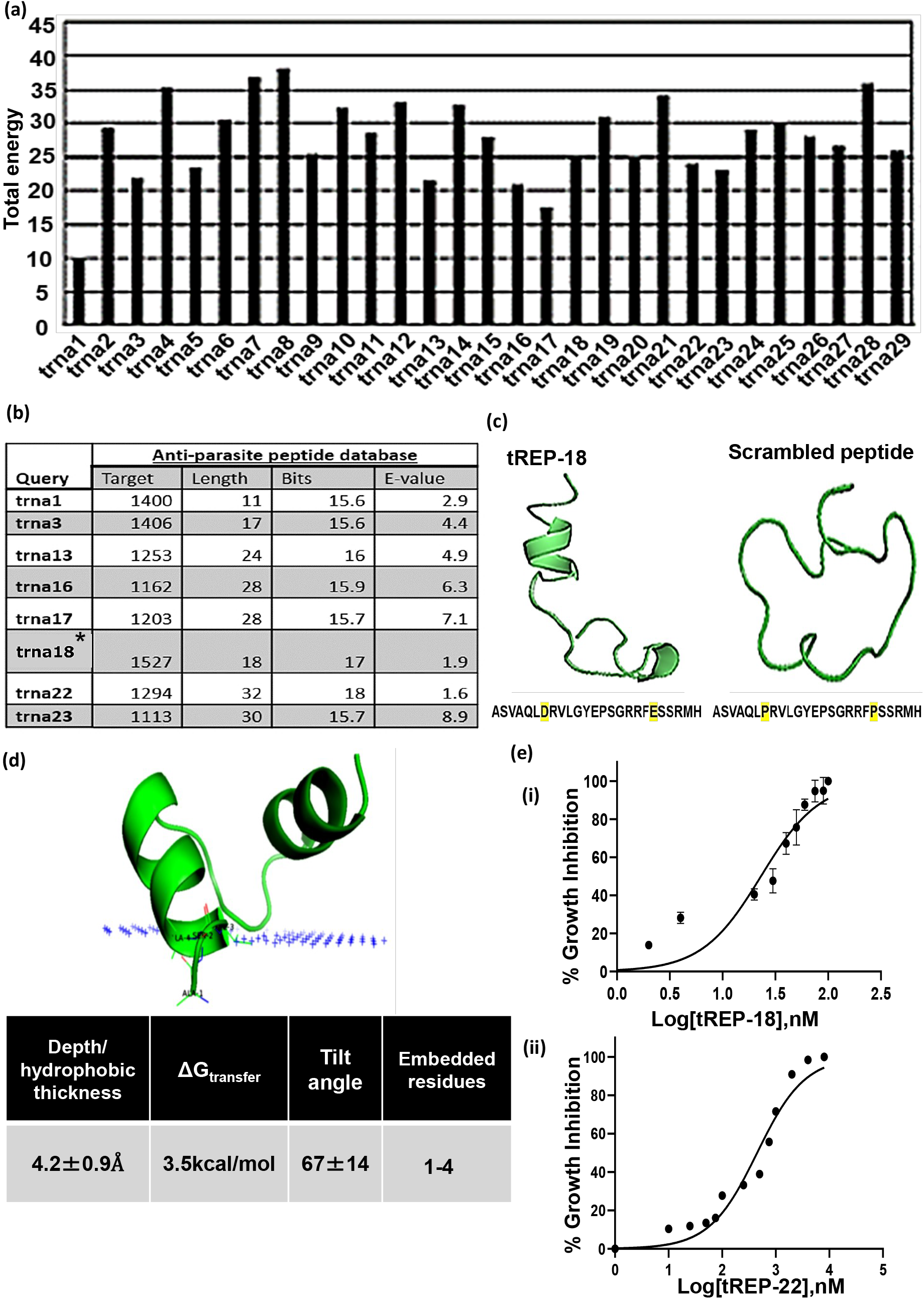
(a) Structural stability analysis of the t-RNA peptides; (b) Structural comparison of tREP18 with scrambled peptide; (c) Sequence similarity analysis against peptide databases; (d) Membrane binding affinity of **tREP18** peptide.

### *In silico* membrane binding studies suggested possibility of tREP18 binding to promastigotes surface

Using Orientations of Proteins in Membranes (OPM) database, we evaluated the binding efficiency of peptide to the membrane of promastigotes *in-silico*. Our results clearly indicated that the peptide is binding with relatively high affinity as evident by the delG energy score = - 3.5 kcal/Mol, and it is embedded into the virtual membrane with a depth of 4.2 Å. The membrane binding affinity of the peptide is found to be moderate, given the minimum depth/hydrophobic thickness in Å. There are 4 residues which are specifically involved in the binding, constituting alanine at 1^st^, 3^rd^ and 4^th^ position; serine at 2^nd^ position (Fig 1d).

### tREP18 is highly cytotoxic and induces cell death in promastigotes

To evaluate the cytotoxic effects of the synthetic peptide on *L. donovani* promastigotes, Lactate Dehydrogenase Assay (LDH assay) was performed. This assay involves reduction of tetrazolium salts to formazan during LDH-mediated catalysis of lactate to pyruvate, which can be detected at 490 nm. Rate of formazan formation is proportional to the release of LDH through damaged cell membranes, which thereby directly correlated with the percentage of dead cells^12^. The promastigotes were incubated with the peptide at various concentrations (1-40 nM) for a period for 24 h, 48 h and 72 h and the released LDH was estimated (Figure 2a). IC50 of **tREP18** is 22.13nM and the CC50 value for the same was found be 310µM and 275µM for HepG2 and J774.2a cells respectively (Supplementary Table 2). Thus, we have used approx. double the concentration of IC50 (40nM) of **tREP18** for all experiments *in vitro*. Amphotericin B treated promastigotes were taken as positive control. The increment in the level of LDH release was observed in both dose and time dependent manner respectively for **tREP18** treatment (Fig 2b). The maximum LDH release was observed in 72 h in amphotericin B treated promastigotes (Avg. O.D. 0.946) representing 100% cytotoxicity. It was observed that the **tREP18** was able to induce the highest level of cytotoxicity (up to 87.86%) at 40nM in comparison to LDH release at 48 h and 24h, which showed 70.27% and 62.21% respectively. Whereas, the scrambled peptide treated samples showed hardly 24.41% cytotoxicity at 72h, thus used as a negative control for further experimental analyses. This result strongly suggests that **tREP18** lead to high level of cytotoxicity as correlated by LDH release. Further to corroborate whether the LDH-based cytotoxicity can be linked to the loss of membrane integrity and cell death, we have monitored propidium iodide (PI) staining in **tREP18**-treated promastigotes [Fig 2c (i)]. The result demonstrated enhancement in PI positivity in a concentration dependent manner, with 95.5% PI positivity upon treatment with 40nM of **tREP18**. This was found to be closer to the level of cytotoxicity induced by Amphotericin B treatment in promastigotes (positive control) showing 97.6% of PI^POS^ cells.

**Figure 2:**
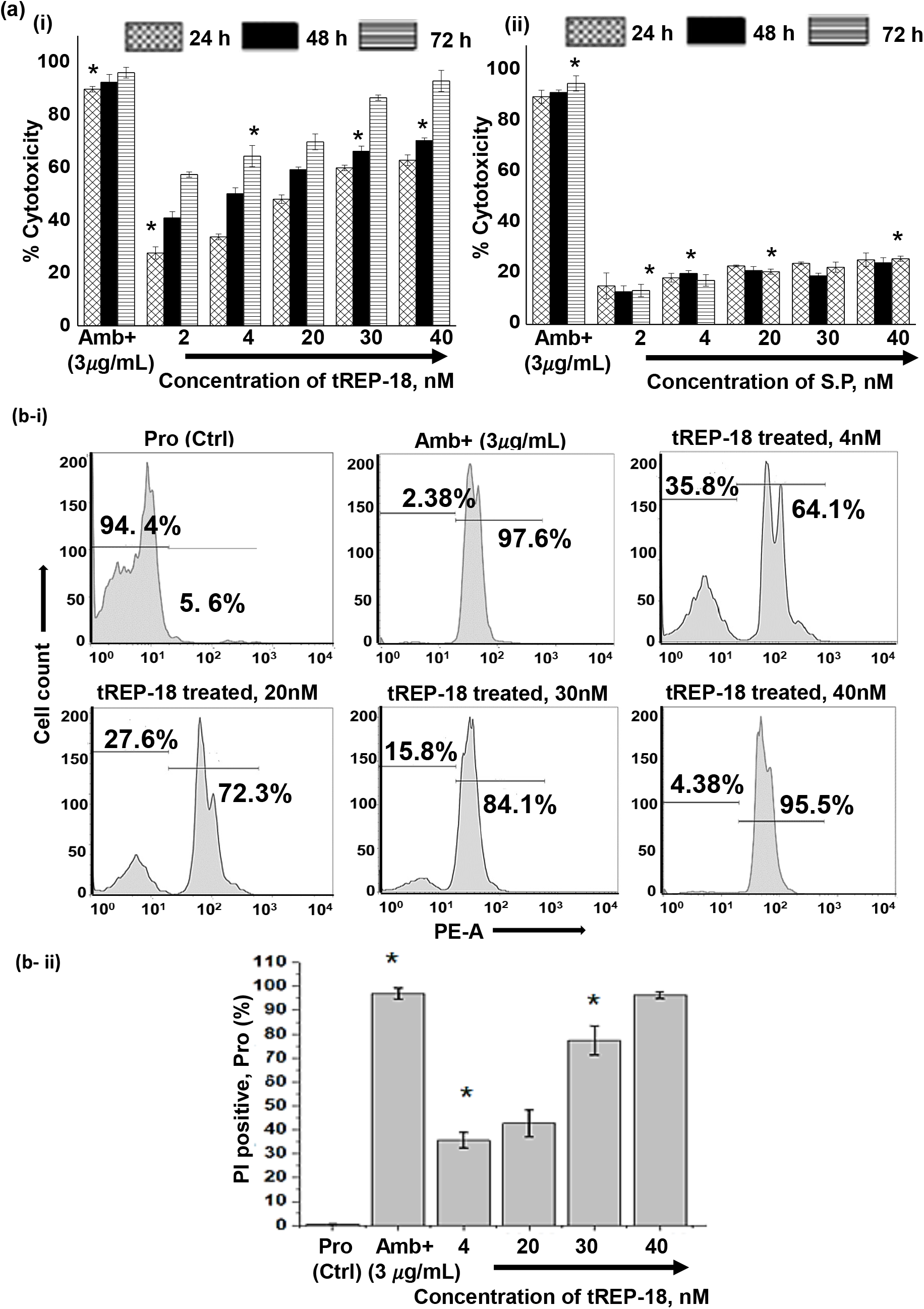
(a) Percentage Inhibition was evaluated using LDH Assay for 24 h following treatments by **tREP18** and scrambled peptide, with 2.5 µg/ml of Amphotericin B was used as a positive control, and its cytotoxic effect (100%) was used for normalization. (b-i) Effect of **tREP18** on promastigotes in a concentration and time dependent manner, manifested as LDH release depicting the cytotoxic effect of novel synthetic peptide; (b-ii) Effect of of scrambled peptide on promastigotes, represented via estimation of LDH release depicting the non-cytotoxic effect of the peptide; (c i) the representative screenshots of PI stained promastigotes at 72 h assessed by flow cytometry analysis depicting parasite death on peptide treatment; (c ii) percentage of PI positivity in **tREP18**-treated promastigotes at 72 h;

### tREP18 caused topographical and morphological alterations in promastigotes

We investigated the morphological and ultrastructural aberrations of the promastigotes following treatment with **tREP18** peptide at 30nM and 40nM respectively using two high-resolution microscopic techniques: scanning electron microscopy (SEM) and atomic force microscopy (AFM). SEM images clearly revealed that at maximum dose of 40nM, the promastigotes lost their self-interacting capability due to rupturing of membrane structure as compared to that of untreated parasites which showed no structural alterations (Fig 3a). Further to understand the ultra-cytoskeletal architecture of cellular phenotype, AFM was employed following treatment with **tREP18** for 72h. Phase scan as well as topographical view of parasites (Fig 3bi) showed profound morphological modulations when treated with 30nM and 40nM concentrations of lead peptide respectively. Analysis of distinctive parameters such as cell length, width and surface roughness prove to be the strong indicators of morphometric integrity. The untreated promastigotes showed typical healthy signatures like elongated spindle-shaped promastigotes with an anterior and long flagellum. Whereas, peptide treated promastigotes demonstrated perturbed membrane, constricted cellular structures and shortened flagella. Ratio of width/length was seen to be lower in untreated control parasites denoting elongated intact structures, whereas **tREP18** treated parasites demonstrated prominently higher width to length ratio suggestive of profound distortion in morphology. Further to determine cell surface roughness, we have estimated the Root Mean Square (RMS value) which is a specific quantitative index to evaluate cell surface topology^13^. The data showed doubling of RMS value (Rq) in treated promastigotes as compared to control ones. Overall, **tREP18** treatment could impose significant alteration in cytoskeletal architecture and surface topology.

**Figure 3:**
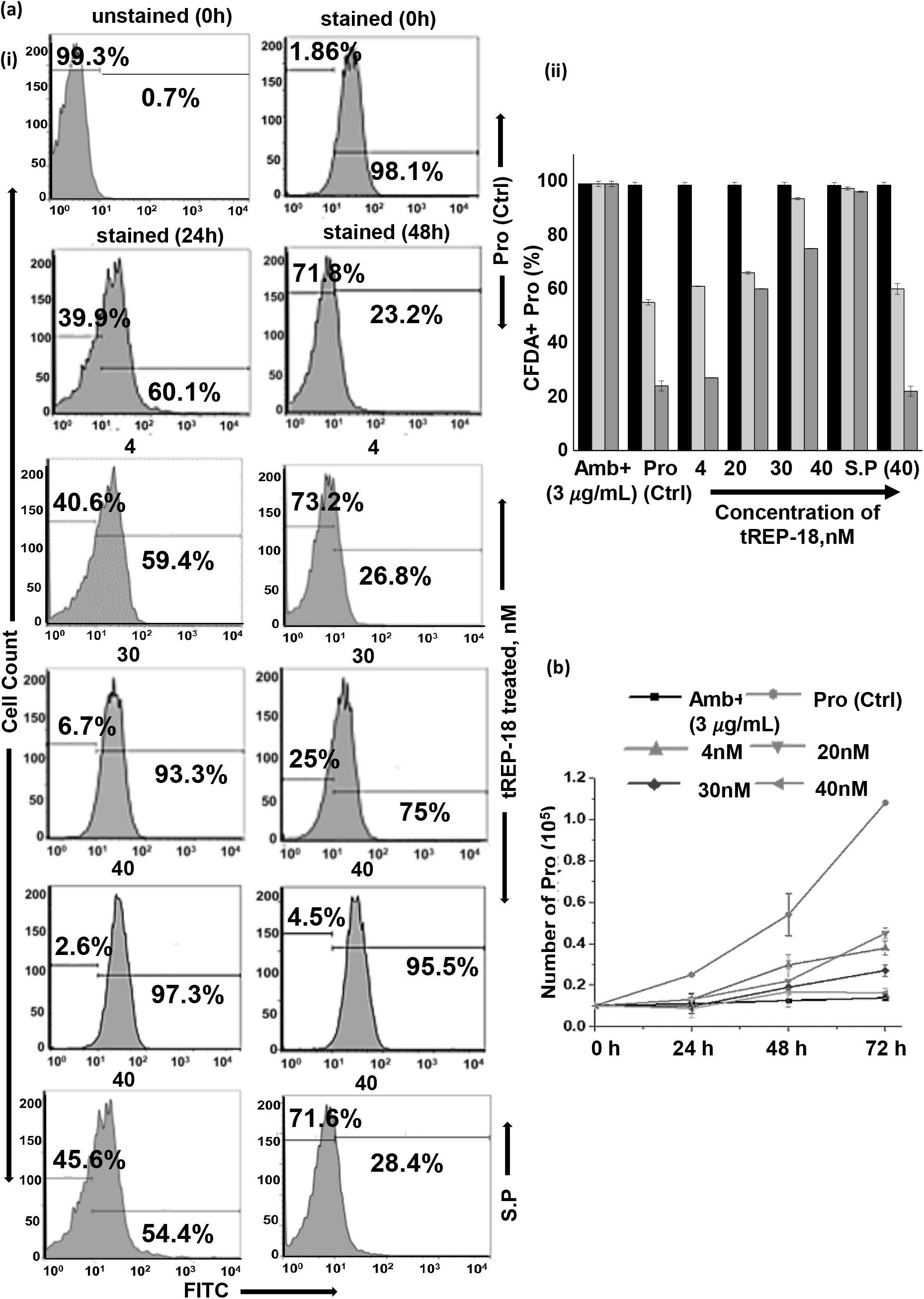
**M**orphological alterations of *Leishmania donovani* promastigotes following treatment with synthetic peptide, (a) Parameters demonstrating surface scanning were elucidated using Scanning Electron Microscopy, (b) parameters showing topological aberrations were depicted using Atomic Force Microscopy,

### tREP18 interferes with mitochondrial function and disrupts mitochondrial membrane potential (ΔΨm)

To explore the effect of **tREP18** on ΔΨm of promastigotes, we have used a lipophilic, cationic dye (JC-1) exhibiting green fluorescence, which enters the mitochondria and gets accumulated into a reversible complex known as J aggregates emitting red fluorescence. The fluorescence intensity was evaluated using flowcytometric analysis and fluorescence microscopy. Thus, the red (PE)/green (FITC) fluorescence ratio of JC-1 in the mitochondria can be considered as a direct assessment of the mitochondria membrane polarization. Enhanced level of red fluorescence denotes more J aggregate formation due to higher ΔΨm, whereas shifting towards lower red or accumulation of higher green fluorescence implies a strong indication of destabilized ΔΨm. Thus, in healthy untreated promastigotes, an intense PE/FITC ratio corresponding to a higher red to green ratio (0.87) was detected representing hyperpolarised mitochondrion, thus suggesting stable ΔΨm (Fig. 4 ai, ii c). However, in case of **tREP18** treated promastigotes, drastic reduction in ΔΨm could be detected with an increasing order of peptide treatment with the maximum effect observed at 40nM, manifested as poor red/green ratio (0.001) exactly similar to amphotericin B treatment (Fig. 4a-ii, c). Based on this result we hypothesize that the disruption of membrane structure lead to compromised ΔΨm on **tREP18** treatment affecting the cellular viability of promastigotes.

**Figure 4:**
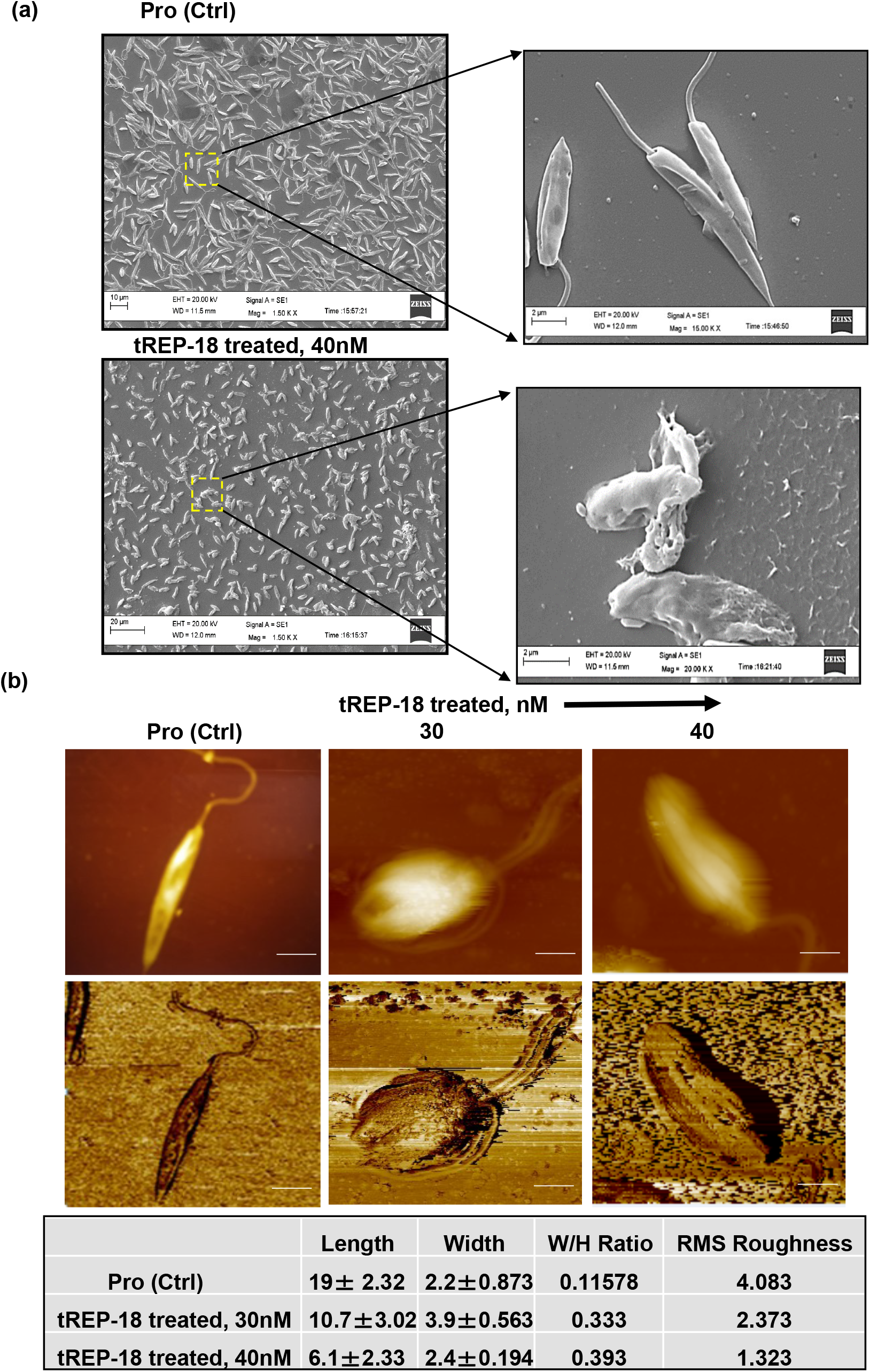
(a & b) Effect of **tREP18** administration on mitochondrial membrane potential (ΔΨm) of promastigotes was determined by the change in JC-1 staining using Flow cytometry and fluorescence microscopy.

### tREP18 affects the growth kinetics and inhibits proliferation of promastigotes

In order to study the impact of **tREP18** on growth dynamics of promastigotes, their replication in culture was monitored for a period of 0-72 h at different treatment conditions (Fig 5a). Parasite density was measured every 24 hours by staining with trypan blue and counting was done in a haemocytometer chamber. An exponential growth curve was observed for untreated cells, while a decreasing pattern of cell growth was evident at 24hours following treatment with increasing concentrations of **tREP18**, including 4nM, 20nM and 30nM respectively. At 72 hours there was 91% decrease in the number of promastigotes in presence of 40nM of **tREP18**, similar to reduction in growth pattern by Amphotericin B treatment, which was used as a positive control (Fig 5a).

**Figure 5:**
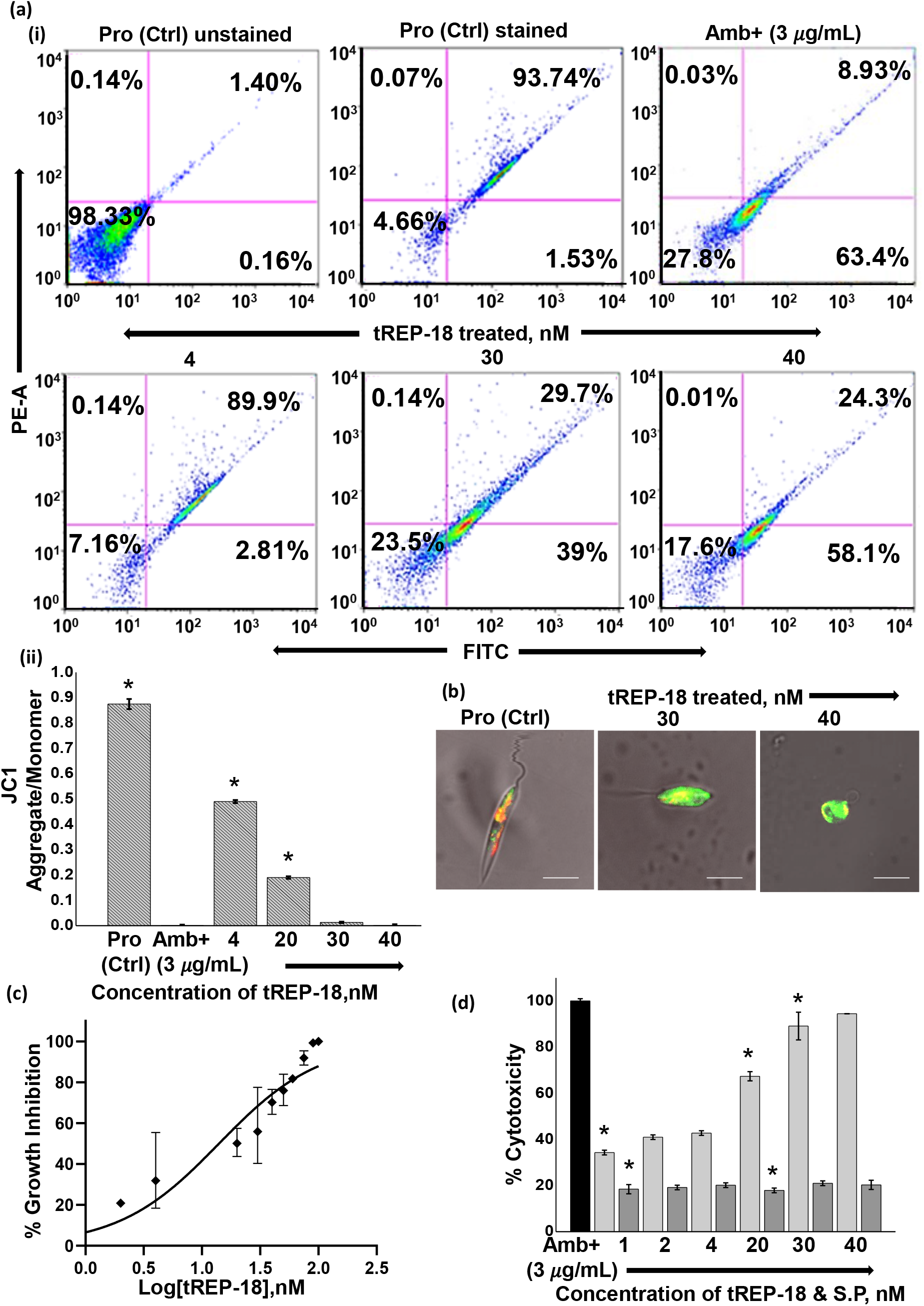
Effect of **tREP18** treatment on promastigotes in dose dependent manner; (a-i) proliferation of daughter cells was determined by reduction in CFDA-SE staining by flow cytometry analysis; (a-ii) percentage of CFDA-SE positivity in **tREP1820**-treated promastigotes; (b) estimation of number of tREP1820-treated promastigotes using trypan blue dye exclusion assay.

To further validate our findings, we assessed the proliferation of promastigotes by quantifying the release of cell permeable dye, CFDA-SE during cell division. This dye enters cells by simple diffusion, following cleavage by intracellular esterase enzymes, to form reactive amine products. Which covalently binds to intracellular lysine residues and other amine sources, producing detectable fluorescence. Decrease in fluorescence intensity is proportional to generation of promastigote daughter cells. The results indicated that, at 40nM concentration of the peptide the percentage of CFDA positive cells were almost remained unchanged at both 24 h (97.3%) and 48 h (96.1%) suggesting no division of the parental cells and indicating no such multiplication in promastigotes number consecutively, as compared to lower concentrations^14^. However, the untreated promastigotes displayed 98.4% of CFDA positivity at 0h, which ultimately reduced to 54.4% at 24 h and 23.2% at 48 h, suggesting cellular progression in a time dependent manner(Fig 5 bi, ii). Taken together, our results demonstrated a significant reduction in promastigotes growth and number on treatment with **tREP18** in a dose dependent manner.

### tREP18 inflicts cytotoxicity in PKDL strain BS12

We further evaluated the effect the **tREP18** and its scrambled control on the clinical isolate of PKDL strain, BS12 of Leishmania sp. The results showed pronounced toxic effect of the **tREP18** on the Indian origin leishmania isolate of PKDL. As mentioned previously, LDH assay was carried out on promastigotes treated with different concentrations of **tREP18** for 72 h, while amphotericin B was used a positive control and scrambled peptide served as negative control. The promastigotes showed minimal toxicity at 1nM with 42% parasitic death and maximum toxicity at 40nM with 98% of cell death like that of amphotericin B treatment (Fig. 6). While, the scrambled peptide showed no toxic effect at all. The data strongly suggests that **tREP18** could adversely affect the metabolic cell viability of clinical strain of PKDL.

**Figure 6:**
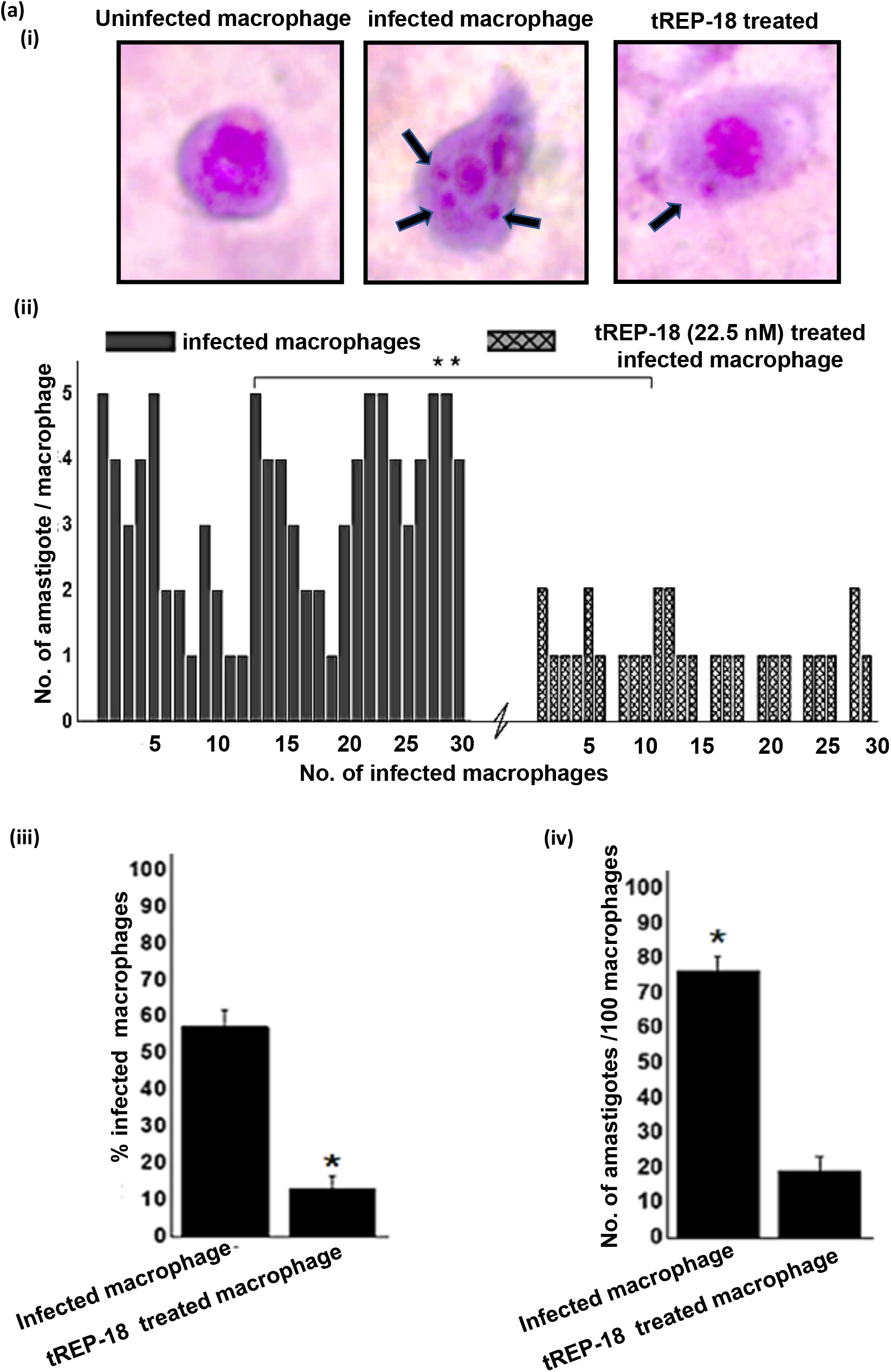
**a)** Cytotoxic effect of **tREP18** peptide and non-toxic effect of scrambled peptide (negative control) on clinical PKDL isolate BS12 respectively.

**Figure. 7.**
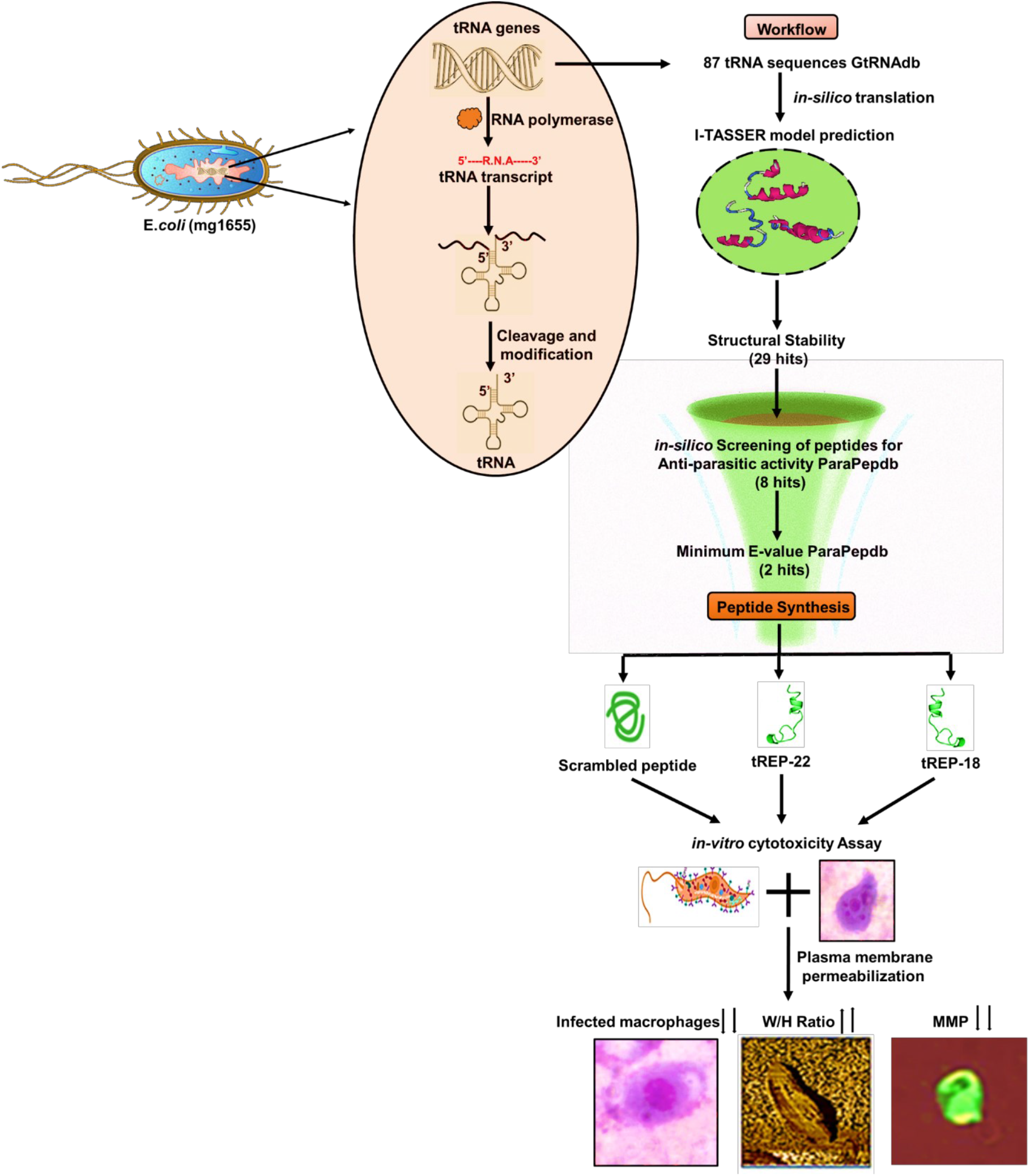

## Discussion

tRNA is an evolutionary invention of non-coding part of genome that needs to be exploited to understand its potential in cellular biogenesis. We hypothesize that, a successful co-ordination between tRNA identification and their translation can lead to birth of novel biomolecular pathways involved in cellular homeostasis. However, the coding potential of tRNA and its possible impact on cell physiology has not been explored yet. Thus, we have addressed the unsolved queries associated with tRNAs involving its possible translation into peptide-like structures and their potential role in biological activities, especially their possible role in targeting eukaryotic pathogen-derived disease, like Leishmaniasis one of the fatal neglected tropical diseases.

To achieve this aim, we have chosen a total of 87 tRNA gene sequences from Escherichia coli genome (mg1655) and translated *in silico* into corresponding peptide sequence. These peptide sequences were chemically synthesized and further used for studying their possible biological functions *in vitro*. Prior to their cellular administration, we have theoretically modelled 3D structures of the peptides, those were energy minimized to identify the most stable ones. The results revealed 9 peptides out of 25 were filtered as the most stable candidate peptides based on e -value <25. Further, only two peptides showed E-value < 2, following in silico screening against anti-parasitic diseases. Out of these two candidates “**tREP18**” with lowest length of 24 mer was considered as the lead peptide, as it is assumed that with lowest length the propensity of proteolytic cleavage lowers down in its cellular targeting. And, with narrow-spectrum activity, **tREP18** might impose no resistance potential along with high selectivity. With these structural attributes, **tREP18** was further used for all experimental studies to evaluate its effect on promastigote form of *L. donovani*. Henceforth, to validate the structural stability and functional importance of **tREP18** peptide, we designed and synthesized a scrambled peptide with change of aspartic acid and glutamic acid at 7^th^ and 20^th^ positions respectively to two prolines, which showed no effect on the promastigotes thus proved to be a negative control.

The cytotoxic effect of the **tREP18** on promastigotes showed a dose dependent increment, and 40nM concentration was chosen as the highest dose for all other experimental evaluations, due to its similar effect as of amphotericin B treatment, that was used as a positive control. Which was also approx. double the IC50 value of **tREP18** (22.13nM) (Supplementary Table 2). We have also screened the CC50 value of **tREP18** in HepG2 and J774.2a cells, which showed nontoxic effects at concentrations > 250µM (Supplementary Table 2). With such functional characteristics, **tREP18** was targeted to *L. donovani* promastigotes to evaluate if it can impose any impact on cytotoxicity in time and dose dependent manner, using lactate dehydrogenase (LDH) as an enzymatic marker of cell death. The finding suggested that **tREP18** can trigger highest release of LDH in a dose and time dependent order with maximum effect at 40nM dose like that of amphotericin B, that further confirmed the lethal effect of our peptide.

The LDH assay results were further corroborated with the PI staining, which also showed that tREP18 could significantly compromise the viability of promastigotes (Fig.2b). To understand if tREP18 treatment could affect the topology of promastigotes, we examined ultrastructural changes using AFM and SEM studies. The result demonstrated that **tREP18** treatment significantly altered the morphology of promastigotes with visible loss in inter-cellular networking and ruptured cellular architecture. Collectively, these results have indicated that **tREP18** induced cell death in promastigotes via severely destabilizing the cellular topology.

Accumulating evidences suggested that collapsed mitochondrial membrane potential is one of the initial events that can lead to cellular death^15^. Based on these, we hypothesized that **tREP18** treatment might lead to depolarization of mitochondrial membrane potential. To determine **tREP18** effect on ΔΨm, we have used a lipophilic cationic dye JC1 monomer that naturally emits green florescence (FITC). In case of healthy cells, due to hyperpolarized state with negatively charged mitochondrial membrane, JC1 can cross the mitochondrial membrane and gets converted to an oligomeric form of reversible J aggregate that emits detectable red fluorescence. Thus, alteration in ΔΨm can be represented as lower red/green ratio suggesting destabilization of the membrane. Our findings revealed that **tREP18** can severely destroy the ΔΨm at 40nM of concentration at 72 h, which is represented by poor red/green ratio (0.001), as compared to control, distorting the redox homeostasis in the cells. It is noteworthy that, destabilization of ΔΨm can lead to generation of reactive oxygen species (ROS), a direct hallmark of apoptosis (). Previous evidences also suggested that enhanced generation of ROS can lead to manifestation of DNA damage, leading to cell death. We assumed that **tREP18** led disruption of ΔΨm is one of the major factors contributing in cell death of promastigotes, which is corroborated by both CFDA and PI positive cell population.

Next, we have evaluated whether treatment of **tREP18** on the promastigotes could also interfere with the growth kinetics of promastigotes using live staining with CFDA-SE, a strong membrane permeant dye. Upon cleavage by esterases within the cell, it generates reactive amine products those covalently bonds with intracellular lysine to make detectable fluorescence products. In case of healthy promastigotes, 98.4% of CFDA positivity could be detected, which gradually reduced to approximately 54.4% and 23% by 24h and 48h respectively, demonstrating a healthy cell division, as clearly indicated by daughter cell generation and dye distribution. Interestingly, **tREP18** treated promastigotes showed no change in dye retentivity at both 24 h (97.3%) and 48 h (96.1%) suggesting, complete abrogation of cell division or cellular progression at all. These experimental outcomes suggest at 40nM concentration **tREP1820** can demonstrate highest efficacy targeting death of *L. donovani* promastigotes.

Based on this background, we then tried to evaluate the efficacy of **tREP18** peptide on clinical PKDL isolate BS12, which is responsible for the post kalazar disease manifestation, and known to play crucial reservoir for *L. donovani* during interepidemic periods. PKDL can impose aggressive manifestation of VL, including devastating facial and skin lesions. Our findings revealed that, **tREP1820** can impart prominent toxic effect even at much lower concentration (1nM), and demonstrated highest toxic effect at 40nM, when kept in contact with promastigotes for a time period of 72 h. Thus, this peptide proves to be a novel therapeutic agent with high efficacy against visceral leishmaniasis caused by *L. donovani* as well as in case of PKDL.

In summary, our study for the first-time reports a novel tRNA-derived peptide with excellent anti-leishmanial property, which can be a potent drug candidate against VL and PKDL.

## Materials and Methods

### Bioinformatics based novel peptide screening

We retrieved the tRNA gene sequences of Escherichia coli strain K-12 sub-strain MG1655, from the genomic tRNA database, that contained tRNA genes. A total of 87 tRNA gene sequences were retrieved and computationally translated into protein sequences using Transeq tool of European Bioinformatics Institute (EBI). Each tRNA sequence was translated into protein sequences and the sequence with stop codon were removed. Further, we computational modeled 3D structure of the peptide using Phypre server. The modeled structures were energy minimized and validated using GROMACS and PROCHECK, respectively. To filter the most stable peptide we subjected the models for FOLDX stability analysis. The stable peptides were screened against the anti-parasite and anti-cancer peptide databases. The peptide with high similarity in both the databases were considered for experimental validation. The scrambled peptide is also modeled using Phypre server.

### Parasite culturing and treatment

Promastigote-form of *Leishmania donovani* (Ag83 strain) was cultured at 26 *C in M199 media (GIBCO, India) pH 7.4 supplemented with 10% (v/v) inactivated Fetal Bovine Serum (GIBCO, India) and 0.02 mg/mL gentamycin (Life Technologies, USA). Cultures were maintained between 10^6^ and 10^7^ cells/mL for continuous exponential growth. 1×10^6^ cells /mL parasite count was constantly maintained for all the experiments. **tREP18** peptide and scrambled peptide were resuspended in dimethyl sulphoxide (DMSO) (Sigma-Aldrich) for preparation of 1M stock solution. 40nM was used as the working concentration for **tREP18** peptide (∼ double the IC50 value) treatment in all *in-vitro* experiments at different time intervals. Parasites without any inhibitor addition were maintained as negative controls.

### Cytotoxicity assay by LDH

LDH cytotoxic assay was performed as per standard protocol (CytoTox 96 Non-Radioactive Cytotoxicity Assay-Promega, USA). Initially, promastigotes were suspended into a 96-well microtiter plate (100 *μ*L well volume). Parasite samples in triplicates were exposed to various concentrations (1nM-40nM) of each peptide and incubated at 26 *C for 72 h. Parasites with Amphotericin B (2.5 *μ*g/mL) (Sigma-Aldrich) was maintained as the positive control. Finally, percentage cytotoxicity of **tREP18** was calculated by normalizing with Amphotericin B treatment that rendered 100% cytotoxicity. Log phase promastigotes (1x 10^6^ /mL) without any peptide addition were maintained as negative controls. Log phase promastigotes (1x 10^6^ /mL) were also treated with scrambled peptide and cytotoxicity of the peptide on parasites was evaluated for 72 h by LDH assay as described in previous experiment.

### Cellular viability and Apoptotic Assay

Promastigotes undergoing apoptosis in both treated and untreated samples were measured by Propidium Iodide (PI) staining. After exposure with the peptide for 24h, 48h and 72h, respectively, cells were harvested, PBS washed and stained with PI (5 *μ*g/mL) (Life Technologies, USA). This was followed by incubation for a period of 20 min at 37°C. Subsequently, cells were washed for excessive stain removal and resuspended in 250 *μ*L PBS. Cells were further analysed through BD FACS diva and also visualized using fluorescence microscope with 510–560 nm filter block for detection of PI red fluorescence.

### Morphological study of promastigotes by Scanning Electron Microscopy

Morphological distortion of **tREP18**-treated promastigotes was examined by SEM. Sample preparation for the SEM was carried out as reported by Gluenz *et. al*. 2012, with slight modification in their protocol. Cells were incubated with 40nM of **tREP1820** for 72 h at 26 °C. These promastigotes were then harvested at 1100g for 15 min at RT and following addition of fresh media. EM-grade glutaraldehyde was directly added to the cells containing M199 media to a final concentration of 2.5% glutaraldehyde (from a 25% stock of EM-grade glutaraldehyde). The cells were centrifuged for 10 mins at 800 g and the media was removed. Promastigotes were then resuspended in 0.1 M phosphate buffer (pH 7.2) and washed twice. These were further fixed with 2.5% (v/v) glutaraldehyde in the same buffer for 120 mins. Glass coverslips were cleaned with ethanol, followed by immersion in a 0.1% (w/v) solution of poly-L-lysine (sigma) in water. Coverslips are then rinsed in water and left to air-dry in laminar hood. 200ul of cell suspension was added onto each coverslip ensuring the coverslip is completely covered by the cell suspension. Coverslips are placed in individual wells of a 12-well tissue culture plate. The plate is left for 10 mins at room temperature for the cells to settle and adhere to the coverslips. Adherence was checked using a tissue culture microscope. Samples were then post fixated in 1% osmium tetroxide for 1 h and dehydrated by gradient acetone concentration (50-100%), 20 min each. Thereafter, samples were treated with 100% hexamethyldisilane at room temperature for 5 min and mounted on aluminium stubs with adhesive carbon tape. Prior to SEM application, a thin gold layer was coated by means of a sputter-coater (SC7640, Polaron Equipment, England, U.K.). The samples were observed under an environmental, variable pressure Scanning Electron Microscope (Carl Zeiss EV0-40, Cambridge, U.K.) at a voltage of 20 kV and a working distance of 10 mm.

### Study of surface topology of tREP1820-treated *L. donovani* promastigotes using Atomic Force Microscopy

Sample preparation for the AFM analysis was carried out as per the protocol of Eaton *et al* 2013. Cells were incubated with 40nM of the peptide for 72 h at 26 °C. These promastigotes were then harvested by centrifugation at 1100g for 15 min at RT, washed with 0.1 M phosphate buffer (pH 7.2) and fixed with 2.5% (v/v) glutaraldehyde in the 0.1M phosphate buffer for 60 min. Cells were washed using the phosphate buffer and overlaid onto poly-L-ornithine (Sigma) coated micro slides having dimension 10mm x10mm. Samples were then washed twice in molecular biology grade water (Sigma) and were dried in laminar hood air flow. Scanning of promastigote cells was carried out with a TT-AFM atomic force microscope. A 50 *μ*m scanner was used, and the instrument operation was done in tapping and non-contact mode. Image details were calculated using XEI software in 1st order flattened 20 x20 µm^2^ areas in the centre of the cell body.

### Quantification of mitochondrial membrane potential

Mitochondrial membrane (Δ*ψ*m) potential was assessed by Fluorescence Assorted Cell Sorting and fluorescence microscopy with 5,6-dichloro-2-[3-(5,6-dichloro-1,3-diethyl-1,3-dihydro-2H-benzimidazol-2-ylidene)-1propenyl]-1,3-diethyl-, iodide (JC-1) (Life Technologies, USA) as a probe. Treated groups and untreated groups were incubated for 24 h. Cells were washed with PBS, JC-1 labelled and samples were analysed through BD FACS diva. The labelled cells were also allowed to adhere to the glass slides for visualization under Fluorescence microscope; excitation and emission filters of TRITC and FITC were used.

### Promastigote Proliferation Assay

Cell growth and multiplication was assessed by Fluorescence Assorted Cell Sorting and fluorescence microscopy with 6-Carboxyfluorescein diacetate succinimidyl ester (CFDA-SE, Life Technologies, USA) as a probe. Promastigotes were washed thrice with 0.1M PBS. The cells were labelled with CFDA-SE dye and were then incubated at 37°C for 10min during which they were mixed 3 to 4 times properly. These cells were then resuspended in ice cold M199 medium. Further they were centrifuged at 1200g for 10 mins (4°C) and resuspended in fresh medium. Cells were treated with the peptide at different concentrations and were analyzed through BD FACS DIVA for 3 consecutive replicates. The labelled cells were also allowed to adhere to glass slides for visualization under Fluorescence microscope; excitation and emission filters of FITC were used.

## Statistical analysis

Student’s t-test was performed to evaluate significant differences between treatment and control samples. P-value < 0.05 was considered to be significant indicated as *. Results represent the mean ± SD of minimum three independent experiments.

## Acknowledgments

This work was supported by funding from DBT builder program, DST, and JNU UPE II program. Dr. Shailja Singh and Dr. Anand Ranganathan are thankful for the funding support from support from Science and Engineering Research Board (SERB, File no. IPA/2020/000007) and Drug and Pharmaceuticals Research Programe (DPRP, Project No. P/569/2016-1/TDT). We sincerely acknowledge help from Prof. Madhubala Rentala, JNU for gifting the J774.A1 murine macrophage cell line. We wish to warmly thank Prof. S. Chandrasegaran, Johns Hopkins University for extremely valuable feedback while reviewing the paper.

## Author contributions

AC, MK, JK and SM performed the experiments, DS, KP and SS performed the bioinformatics study, AR, JS and AR helped in experiments, SP and SS (corresponding author) mentored and provided experimental support. PKD conceived the idea, designed the project, analysed the results and wrote major part of the publication.

